# An antibody-drug conjugate targeting GPR56 demonstrates efficacy in preclinical models of colorectal cancer

**DOI:** 10.1101/2022.08.11.503187

**Authors:** Joan Jacob, Liezl E. Francisco, Treena Chatterjee, Zhengdong Liang, Shraddha Subramanian, Qingyun J. Liu, Julie H. Rowe, Kendra S. Carmon

## Abstract

Antibody-drug conjugates (ADCs) have become an increasingly successful class of anticancer therapy, particularly within the past few years. Though ADCs are in clinical trials for colorectal cancer (CRC), a candidate has yet to be approved. CRC continues to be a leading cause of cancer-related death, emphasizing the need to identify novel target antigens for ADC development. GPR56, a member of the 7TM receptor family, is upregulated in colorectal tumors compared to normal tissues and located on the surface of CRC cells, making it a promising ADC target. Furthermore, high GPR56 expression occurs in tumors that are microsatellite stable, negative for the CpG methylator phenotype, and show chromosomal instability. We previously reported the generation of a high affinity GPR56-specific monoclonal antibody, 10C7, and we have now mapped the epitope to the C-terminal end of the extracellular domain, proximal to the GPCR proteolysis site. Here, we describe the development of a duocarmycin-conjugated 10C7 ADC. 10C7 co-internalized with GPR56 and trafficked to the lysosomes of CRC cells, which is critical for efficient ADC payload release. Evaluation of the ADC in a panel of CRC cell lines and tumor organoids with different levels of GPR56 expression showed the ADC selectively induced cytotoxicity at low nanomolar concentrations in a GPR56-dependent manner. A nontargeting control ADC showed minimal to no activity. Furthermore, GPR56 ADC exhibited significant antitumor efficacy against GPR56-expressing patient-derived xenograft models of CRC. This study provides rationale for the development of a GPR56-targeted ADC approach to potentially treat a large fraction of CRC patients.

## Introduction

Colorectal cancer (CRC) continues to be a leading cause of cancer-related morbidity and mortality worldwide (1). Standard of care treatment is most commonly surgical resection followed by chemotherapy and/or radiation therapy. Long-term prognosis remains meagre for patients with metastatic and recurrent disease. Chemotherapy has been limited by systemic toxicity, low response rates, and drug resistance. Targeted therapies such as anti-EGFR monoclonal antibodies (mAbs) have improved clinical outcomes, yet only a fraction of patients with wild-type *KRAS/NRAS/BRAF* benefit (2). More recently, immune checkpoint inhibitors have shown to be effective in a subset of patients with microsatellite instability-high (MSI-H) metastatic CRC (2,3). Given that the majority of CRC patients (80-85%) are classified as non-MSI-H or microsatellite stable (MSS), indicates there is still an urgent need for new targeted therapies and approaches to CRC treatment.

Antibody-drug conjugates (ADCs) are becoming increasingly important therapeutics in the cancer treatment landscape. To date, 12 ADCs have been approved in the United States for the treatment of hematological malignancies and solid tumors (4), with 7 receiving approval in just the past few years. This weaponized antibody therapy is designed for selective delivery of highly potent payloads to tumor cells by linking them to monoclonal antibodies, thereby enhancing the therapeutic efficacy while minimizing systemic effects (5,6). ADCs are composed of three main components that can influence the therapeutic index: the antibody, linker, and payload. Typical payloads being used in ADC development have diverse mechanisms of action, including microtubule inhibitors (e.g. auristatins and maytansinoids), topoisomerase inhibitors (e.g. camptothecin, exatecan), and DNA-damaging agents (e.g. duocarmycin and pyrrolobenzodiazepines) (7). As ADC technology continues to evolve and expands to include novel targets and linker-payloads, it persists as one of the fastest growing classes of anticancer therapy.

Here we describe the preclinical development of a unique GPR56-targeted ADC for the treatment of CRC. GPR56 is a member of the adhesion G protein-coupled receptor (GPCR) subfamily, and is also known as ADGRG1 (adhesion GPCR subtype G1). It is highly expressed in a large fraction of CRC and other tumor types (8–11), with lower expression in normal tissues (10,12–15). GPR56 correlates with poor prognosis and survival (11). We and others showed that GPR56 is associated with drug resistance and promotes tumor growth in mouse models of CRC (11,16,17). Further analysis of RNAseq datasets and protein expression shows that high GPR56 expression is associated with non-MSI-H, non-CIMP (CpG island methylator phenotype), and chromosomal instability (CIN+). Previously, we generated 10C7, a GPR56-specific mAb that binds with high affinity and potentiates GPR56-mediated signaling in vitro (18). In this study, 10C7 was used for the development of a GPR56 ADC, conjugated to the DNA minor-groove-binding, alkylating payload, duocarmycin, via a protease-cleavable linker. The GPR56 ADC showed high selectivity and potency against GPR56-high CRC cell lines and antitumor efficacy in patient-derived xenograft models. This study demonstrates a novel GPR56-targeted approach to potentially treat a large fraction of non-MSI-H colorectal tumors.

## Methods and Materials

### Plasmids and cloning

Reporter vector pGL4.34[luc2P/SRF-RE/Hygro] was purchased from Promega. The myc-tagged hGPR56 plasmid (pIRESpuro3-myc-hGPR56) was previously described (11) and human to mouse GPR56 mutants were generated by site-directed mutagenesis using standard procedures and the primers listed in (Supplemental Table 1). Mouse Gpr56 cDNA (CloneId:3709247) was purchased from Horizon Discovery and myc-tagged mGPR56 was cloned as described for hGPR56. All expression constructs were sequence verified.

### Commercial antibodies and payloads

Commercial primary antibodies used in this study include anti-myc-tag (2276), anti-β-actin (4970), and anti-LAMP1 (9091), all from Cell Signaling. Free ADC payload DM1 was purchased from MedChemExpress, DX8951 from Levena, MMAE from ALB Technology, and Duocarmycin SA from The Chemistry Research Solution LLC.

### Antibody production and antibody-drug conjugate generation

Anti-GPR56 mAb 10C7 was generated and sequenced as previously described (18). For large-scale production, plasmids containing the light and heavy chain of 10C7 or a non-targeted control hIgG1 (cmAb) were transiently expressed in Expi293F suspension cells (ThermoFisher) using polyethylenimine reagent. Mab-containing medium was harvested approximately 7 days post-transfection. MAbs were purified using CaptivA^®^ Protein A Affinity Resin (Repligen), for affinity chromatography, and then eluted in PBS. GPR56 ADC and non-targeting isotype control ADC (cADC) were generated by Levena Biopharma using a cysteine-based conjugation as we have previously described (19). Briefly, partial reduction of the interchain disulfide bonds of the mAbs was performed to generate reactive cysteine thiol groups, followed by conjugation of the linker-payload, mc-vc-PAB-DMEA(PEG2)-Duocarmycin SA. ADCs and mAbs were analyzed by Coomassie blue stained SDS-PAGE for purity, size exclusion chromatography (SEC-HPLC) to measure aggregation and hydrophobic interaction chromatography (HIC-HPLC) to determine drug-antibody ratio (DAR). ADCs had ≥ 98% monomers by SEC-HPLC (i.e. less than 2% high molecular weight aggregates and/or fragments). An average DAR of 3.54 and 3.9 was determined for GPR56 ADC and cADC, respectively.

### Cell Culture and transfection

DLD1, RKO, SW48, LS180, HCT116, COLO320, HCT15, HT29, SW403, SW620, LoVo and HEK293T (293T) cells were purchased from ATCC. Cell lines were authenticated utilizing short tandem repeat profiling and routinely tested for mycoplasma. 293T cells were cultured in DMEM and CRC cell lines in RPMI medium supplemented with 10% fetal bovine serum and penicillin/streptomycin. Transient transfections were performed using jetPRIME (Polypus Transfection). Mutant H360H and mGPR56 293T stable cell lines were established as described for vector and hGPR56 stable lines (18). DLD-1 CTL and GPR56 KD (shRNA) stable cell lines were established as previously reported (11).

### Patient-derived tumor organoids

The colorectal tumor organoid model was established from a liver metastasis of rectal adenocarcinoma harvested from a tumor-bearing patient-derived xenograft (PDX) mouse (TM00849, The Jackson Laboratory). To generate organoids, tumor tissue was cut into small pieces, washed with ice-cold PBS, and subsequently digested with Liberase (TH grade; Roche Life Science) for 1 hr at 37°C with vigorous pipetting every 15 min. The remaining fragments were treated with TrypLE Express (Invitrogen) at 37°C for 20 min to disperse into single cells. The supernatant was collected and centrifuged at 200 g for 3 min at 4°C. The cell pellet was suspended with growth factor reduced Matrigel (Corning) and dispensed into 48-well culture plates (25 μl Matrigel/well). After Matrigel polymerization, complete medium (Advanced DMEM/F12 supplemented with 50 ng/ml EGF, 100 ng/ml Noggin, 500 nM A83-01, 10 nM Gastrin, 1x B-27, 1 mM N-acetylcysteine, 10mM HEPES, 2mM Glutamax, and penicillin/streptomycin) was added and replenished every 2-3 days.

### Western blot analysis

For western blots, protein extraction was performed using RIPA buffer (Sigma) supplemented with protease/phosphatase inhibitors. Cell lysates were incubated at 37°C for 1 hr and diluted in laemmli sample buffer prior to loading on SDS-PAGE. Commercial antibodies were used in accordance to the manufacturer’s guidelines and HRP-labeled secondary antibodies were utilized for detection with the standard ECL protocol.

### Antibody internalization

For immunocytochemistry (ICC), CRC cell lines DLD-1, HCT15, SW403, LoVo or vector and hGPR56 293T cells were seeded into 8-well chamber slides (BD Biosciences) and incubated overnight. The next day, cells were treated with 10C7 or anti-myc-tag mAb at 37°C for 30 min or 1 hr in 293T cells (as indicated) or 1 hr in CRC cells to allow binding and internalization of mAbs. Cells were washed with PBS, fixed in 4% formalin, permeabilized in 0.1% saponin, then incubated with anti-LAMP1 for 45 min at room temperature for lysosome colocalization studies. Cells were then incubated with corresponding secondary antibodies for 1 hr at room temperature: goat anti-human-Alexa-488 and goat anti-mouse-Alexa-488 (Invitrogen) for 293T internalization studies and goat anti-rabbit-Alexa-488 and goat anti-human-Alexa-555 (Invitrogen) for lysosome colocalization studies. Nuclei were counterstained with TO-PRO-3. Images were acquired using confocal microscopy (Leica TCS SP5 microscope) with the LAS AF Lite software (Leica Microsystems, Inc.).

### Cell-based binding assays

For whole-cell based binding assays, stable 293T cells overexpressing vector, hGPR56, mGPR56 or H360S mutant were seeded onto poly-D-lysine-coated 96-well plates and incubated overnight. Serial dilutions of 10C7 or GPR56 ADC were added for 2 hrs at 4°C. Plates were washed in PBS, fixed in 4% formalin, and incubated with goat anti-human-Alexa-555 (Life Technologies) for 1 hr at room temperature. Plates were washed with PBS and fluorescence intensity was quantified using Tecan Infinite M1000 plate reader. Data were analyzed and Kd values were determined using Prism (GraphPad Software, Inc.).

### Luciferase-reporter assays

Reporter assays were performed as previously described (18). Briefly, 293T cells were transiently transfected in 6-well plates with pGL4.34 (SRF-RE) and other expression vectors, as indicated. The next day, cells were plated at 2500 cells/well in 96 half-well plates. Serial dilutions of 10C7 was added and incubated overnight at 37 °C. Luciferase activity was measured using Pierce Firefly Luciferase Glow Assay Kit according to manufacturer’s protocol using an EnVision multilabel plate reader (PerkinElmer). AlamarBlue (ThermoFisher) was used for normalization to cell viability and fluorescence quantified at 530/590 nm using Tecan Infinite M1000 plate reader. Each condition was performed at least 3 times in triplicate.

### In vitro cytotoxicity

CRC cells were plated at 1000-1500 cells/well and 293T cells plated at 500 cells/well in 96 half-well plates. Serial dilutions of unconjugated GPR56 mAb, cmAb, GPR56 ADC or cADC were added and incubate at 37°C for 3 (293T) or 4 days. For tumor organoid viability assay, 1000 cells/well (passage 3) were seeded into a 48-well plate and after 3 days, media was replaced with fresh medium before treatment. GPR56 ADC or cADC (30nmol/L) was added to cells or organoids and allowed to incubate at 37°C for 5 days. Viability was measured using CellTiter-Glo^®^ assay (Promega) for cell lines or CellTiter-Glo^®^ 3D assay for tumor organoids, according to the manufacturer’s protocol. Luminescence was measured using EnVision mulitlabel plate reader (PerkinElmer). Bright-field images of organoids were acquired using an Olympus IX71 microscope. Representative results of at least 3 independent experiments are shown. IC_50_s were determined using Prism (GraphPad Software, Inc.).

### In vivo efficacy studies

Female nu/nu (Charles River or Jackson Laboratory) and NSG (The Jackson Laboratory) mice were used for CRC cell lines and patient-derived xenograft models, respectively. The patient-derived xenograft CRC-001 model was purchased from Jackson Laboratory (TM00849) and the XST-GI-005 model was established in our laboratory at UTHealth with patient consent and appropriate approvals (HSC-MS-20-0327, HSC-MS-21-0074, AWC-20-0144). Female 6-8-week-old mice were implanted subcutaneously with 10^6^ CRC cells (SW620, HT-29, SW403) or 2-3 mm tumor fragments (CRC-001 and XST-GI-005) into the right flank. When tumors reached approximately ~100-200 mm^3^, mice were randomized into treatment groups. Mice were intravenously dosed weekly with vehicle (PBS), GPR56 mAb, cADC, or GPR56 ADC as indicated. Mice were routinely monitored for morbidity and mortality. Tumor volumes were measured bi-weekly and estimated by the formula: tumor volume = (length x width^2^)/2. Mice were euthanized when tumor diameter reached 15 mm.

### Statistical Analysis

All data were analyzed using GraphPad Prism software. Data are presented as mean ± SEM or ± SD, as indicated. Statistical significance of in vitro experiments was analyzed with one-way analysis of variance (ANOVA) and Tukey’s multiple comparison test. Statistical analysis of tumor volume was assessed by oneway ANOVA and Dunnett’s multiple comparison test or Student’s t test for two groups. P value less than < 0.05 was considered significant.

## Results

### GPR56 is highly expressed in the non-MSI-H and non-CIMP colorectal cancer molecular subtypes

We previously showed that GPR56 is upregulated in colorectal tumors compared to matched normal tissues from different cohorts and its expression correlates with poor disease-free and overall survival (11). To further identify the patient group that would most likely benefit from an GPR56-targeted ADC therapy, we evaluated GPR56 association with molecular subtypes. TCGA colorectal adenocarcinoma (COADREAD) dataset consisting of a total of 50 normal and 383 tumor samples showed *GPR56* mRNA is significantly higher in tumors (average fold difference ≈ 2.7; Fig. 1A)(12,20). We next analyzed 243 tumor samples (21), including 35 MSI-H (14.4%), 42 MSI-L (17.3%) and 166 MSS (68.3%). The *GPR56* expression levels were significantly upregulated in MSI-L and MSS tumors compared with MSI-H tumors (Fig. 1B). Of note, MSI-L is typically grouped with those tumors classified as MSS. As MSS tumors typically present chromosomal instability (CIN), we also showed that *GPR56* mRNA is higher in CIN+ tumors (Fig. 1C). The COADREAD study identified four methylation clusters, namely CpG island methylator phenotype-high (CIMP-H), -low (CIMP-L), Cluster 3, and Cluster 4, based on unsupervised clustering of promoter DNA methylation profiles (21). For this analysis, Cluster 3 and Cluster 4 were designated as non-CIMP. As shown in Fig. 1D, *GPR56* expression levels were significantly increased in CIMP-L and non-CIMP compared to CIMP-H tumors. *GPR56* levels were higher in non-CIMP compared to CIMP-L, however the difference was not significant. We next investigated if *GPR56* expression correlated with mutational statuses of *KRAS, TP53*, or *BRAF* (Fig. 1F-G). No association with *KRAS* or *TP53* was identified. However, GPR56 was negatively associated with mutation in *BRAF*, as this mutation typically occurs in MSI-H tumors (21). CRC RNA-seq data from the Cancer Cell Line Encyclopedia was consistent with COADREAD data showing higher levels of *GPR56* in MSS and non-CIMP subtypes, though the latter was not significant (Supplementary Fig. S1A-B). Classification of CRC cell lines based on molecular subtypes was previously reported (22,23). Western blot analysis of a panel of CRC cell lines showed GPR56 protein expression was highest in MSS cell lines HT-29, SW403, and SW620, the latter two of which are also non-CIMP (Fig. 1H). MSS cell line Colo320 had no detectable expression. For MSI-H cell lines, moderate expression was detected in DLD-1, LS180, and HCT15 and minimal to no expression was detected in LoVo, RKO, SW48, and HCT116. These results show that GPR56 is most highly expressed in patient tumors of the non-MSI-H, non-CIMP, CIN+ molecular subtype.

**Figure 1.**
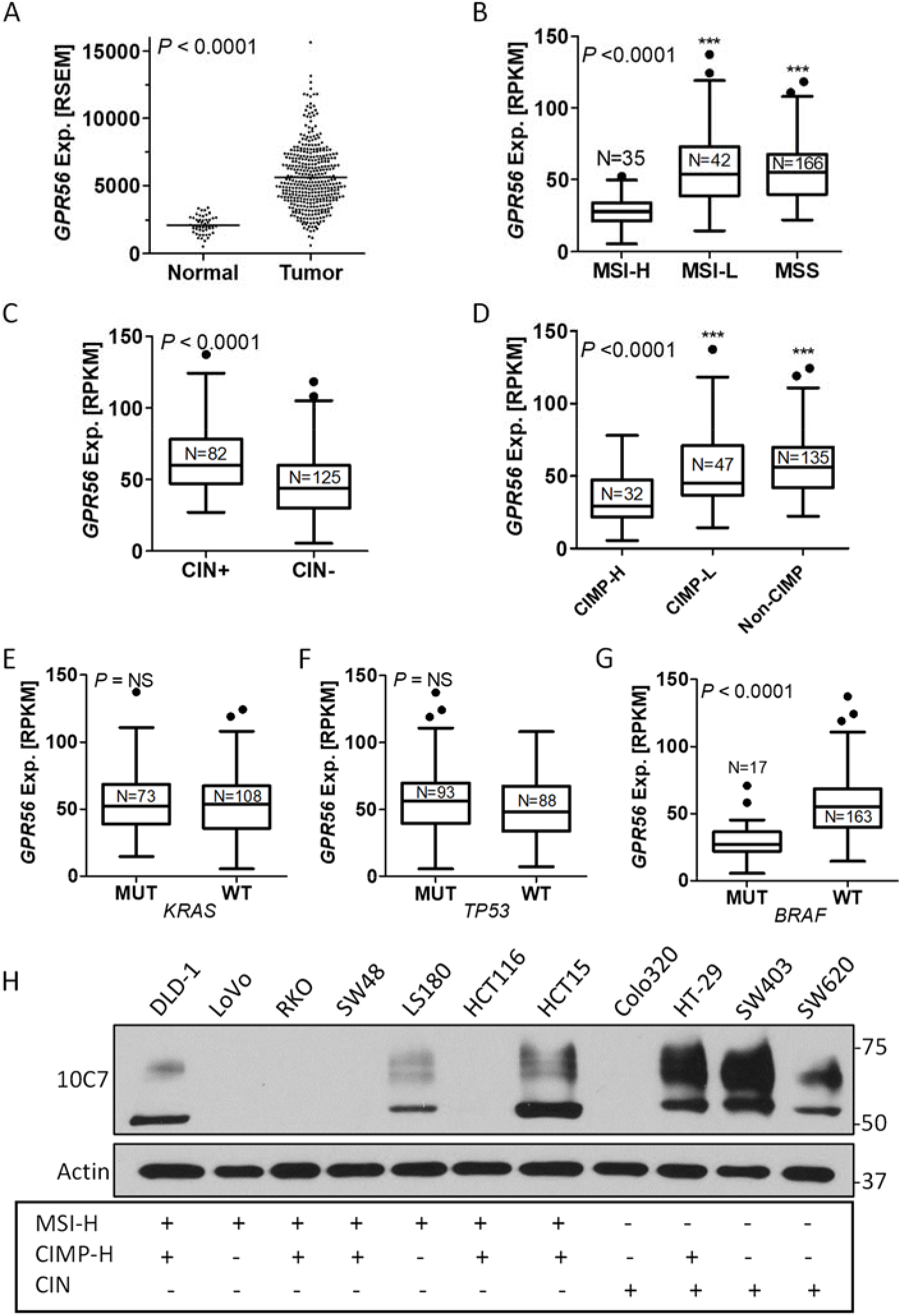
GPR56 expression and correlations with microsatellite instability (MSI), chromosomal instability (CIN), CpG island methylator phenotype (CIMP), and mutational statuses in colorectal cancer. Box-and-whiskers plots showing A, GPR56 RNAseq expression data of tumor (N=383) and normal tissues (N=50) from the TCGA colorectal adenocarcinoma (COADREAD) cohort. RSEM, RNA-Seq by expectation-maximization. Distribution of B, MSI/MSS, C, CIN and D, CIMP subtypes based on GPR56 expression analysis. GPR56 expression levels in tumors with WT and mutations in E, KRAS F, TP53 and G, BRAF. RPKM, reads per kilobase of transcript per million. H, Western blot of GPR56 protein expression in a panel of colorectal cancer cell lines of different molecular subtypes. Statistical analysis performed using one-way analysis of variance (ANOVA) and Tukey’s multiple comparison test or Student’s t-test for two groups.

### 10C7 mAb binds the GPR56 extracellular domain proximal to the GPCR proteolysis site (GPS)

We previously reported the generation of 10C7, a mAb that binds hGPR56 with high affinity (Kd ≈7-8 nmol/L) and can potentiate GPR56-mediated serum response factor-response element (SRF-RE) activity (EC_50_ ≈2 nmol/L; (18)). Studies using deletion mutants showed that 10C7 binds within the region of the G-protein-coupled receptor (GPCR) autoproteolysis-inducing (GAIN) domain, spanning amino acid (AA) residues M176 to the GPCR proteolysis site (GPS) between L382 and T383 (Fig. 2A; (18)). To further characterize 10C7 prior to ADC generation, we set out to evaluate species specificity and map the AA residue(s) important for GPR56 binding and activation. Using a cell-based fluorescence binding assay, we showed 10C7 had high-affinity binding for hGPR56 with no binding to vector 293T cells, as previously reported (18). However, 10C7 failed to bind 293T stable cells overexpressing mouse GPR56 (mGPR56) (Fig. 2B). Given the sequence similarity between human and mouse GPR56 in this region is only ~78% (162/208 aa), we performed a human to mouse site-directed mutagenesis screen to pinpoint the residue(s) critical for 10C7 binding (Fig. 2A). Western blot analyses showed that 10C7 was able to detect expression of all hGPR56 mutants except H360S (Fig. 2C and Supplementary Fig. S2A). As WT and mutants all express a myc-tag at the N-terminus, a myc-tag mAb was used to confirm protein expression (Fig. 2C). Since 10C7 was still able to detect P358T and R370S, E372D mutants (Fig. 2C and Supplementary Fig. S2A), this suggests that the epitope region lies between aa G359 and V369, and contains H360. To further verify that H360 is a critical residue of the 10C7 epitope, we performed a cell-based fluorescence binding assay using 293T stable cell lines overexpressing hGPR56 WT or H360S and quantified binding. 10C7 showed similar, high-affinity binding for WT, as previously reported (Carmon JBC). No binding of 10C7 was detected in H360S or vector 293T cells (Fig. 2D). Immunocytochemistry (ICC) and confocal analysis confirmed both 10C7 and myc-tag mAbs bind to WT cells, whereas only the myc-tag mAb displayed binding to H360S cells (Fig. 2E). No binding was detected for either mAb in vector cells, as expected (Fig. 2D-E). We next tested the effect of H360S mutation on 10C7-mediated activation of GPR56-mediated signaling using the SRF-RE reporter assay in 293T cells. Of note, this assay measures transcription of a luciferase reporter gene in response to G12/13-RhoA-mediated activation of SRF. Transient overexpression of WT or H360S similarly increased SRF-RE basal activity compared to vector (Fig. 2F). WT showed a dosedependent increase in SRF-RE activity in response to 10C7, however H360S and vector activity was unchanged. Equivalent expression of WT and H360S was confirmed by western blot (Supplementary Fig. S2B). Together, these findings indicate that residue H360, proximal to the GPS site, is a critical residue for 10C7 binding and activation of hGPR56.

**Figure 2.**
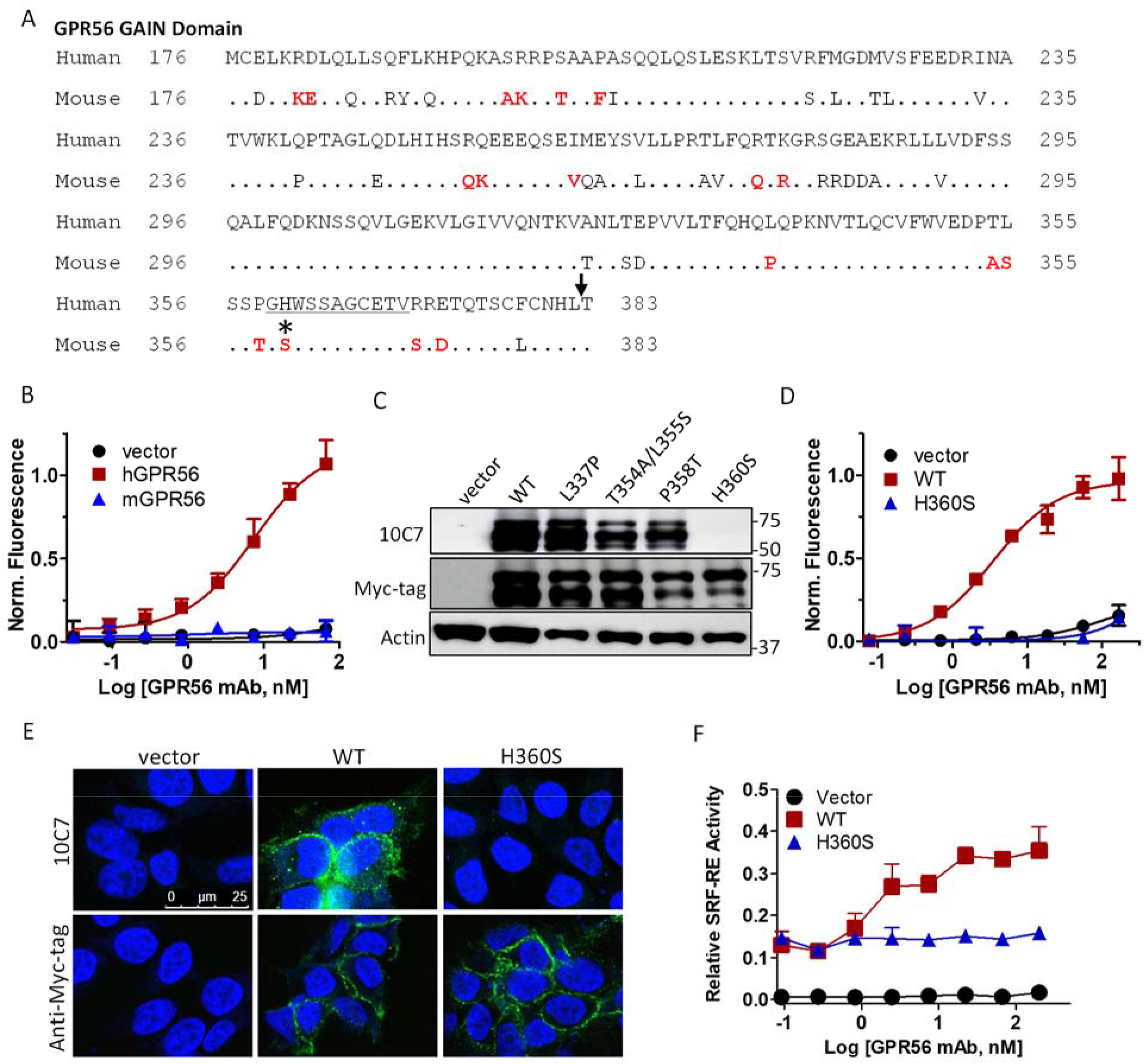
Mapping of the 10C7 mAb epitope within the GAIN domain of the hGPR56 ECD. A, Sequence alignment of amino acids 176-383 of the GAIN domains of human and mouse GPR56. Amino acids in red indicate where mutations were made to convert the human to mouse sequence for epitope mapping. Arrow indicates GPCR proteolysis site (GPS). *, represents histidine critical for mAb binding. B, Cell-based binding assay shows 10C7 mAb binds hGPR56-overexpressing 293T cells and not vector or mGPR56 cells. C, Western blot of 10C7 detection of different hGPR56 mutants. D, Cell-based binding assay and E, confocal microscopy images show hGPR56 H360S mutation inhibits 10C7 binding. For ICC, 293T cells were treated with 10C7 (30 nmol/L) or myc-tag mAb for 30 min at 37 °C. F, SRF-RE luciferase reporter assay shows 10C7 activates WT, but not H360S mutant signaling in a concentration-dependent manner. Experiments were performed 3 times. Error Bars are SD.

### 10C7 co-internalizes with GPR56 and traffics to the lysosome

To make an effective GPR56 ADC, 10C7 must bind endogenous GPR56 on live cells, internalize, and be delivered to the lysosome for payload release. As the 10C7 epitope is not completely conserved between human and mouse, a 10C7-based ADC can only be tested for proof-of-concept against human models. To evaluate internalization and intracellular trafficking, vector and hGPR56 293T cells were incubated with 10C7 for 1 hour at 37°C. ICC and confocal microscopy showed that 10C7 (red) effectively internalized and co-localized with the lysosome marker LAMP1 (green) in hGPR56 293T cells (Fig. 3A). Co-localization is indicated by yellow signal in the merged image. No binding was detected in control vector cells. We next evaluated 10C7 binding and internalization with endogenous GPR56 using several CRC cell lines with different levels of GPR56 expression. After a 1-hour incubation, we observed strong intracellular staining of 10C7 and colocalization within lysosomes in DLD-1, HCT15, and SW403 cells (Fig. 3B). No binding or internalization signal was observed in DLD-1 cells with a nontargeted hIgG1 control antibody (cmAb) or in GPR56-negative LoVo cells with 10C7 (Fig. 3B, top and bottom panels). These data show that 10C7 binds endogenous GPR56 with high specificity and internalizes to the lysosomes of CRC cells.

**Figure 3.**
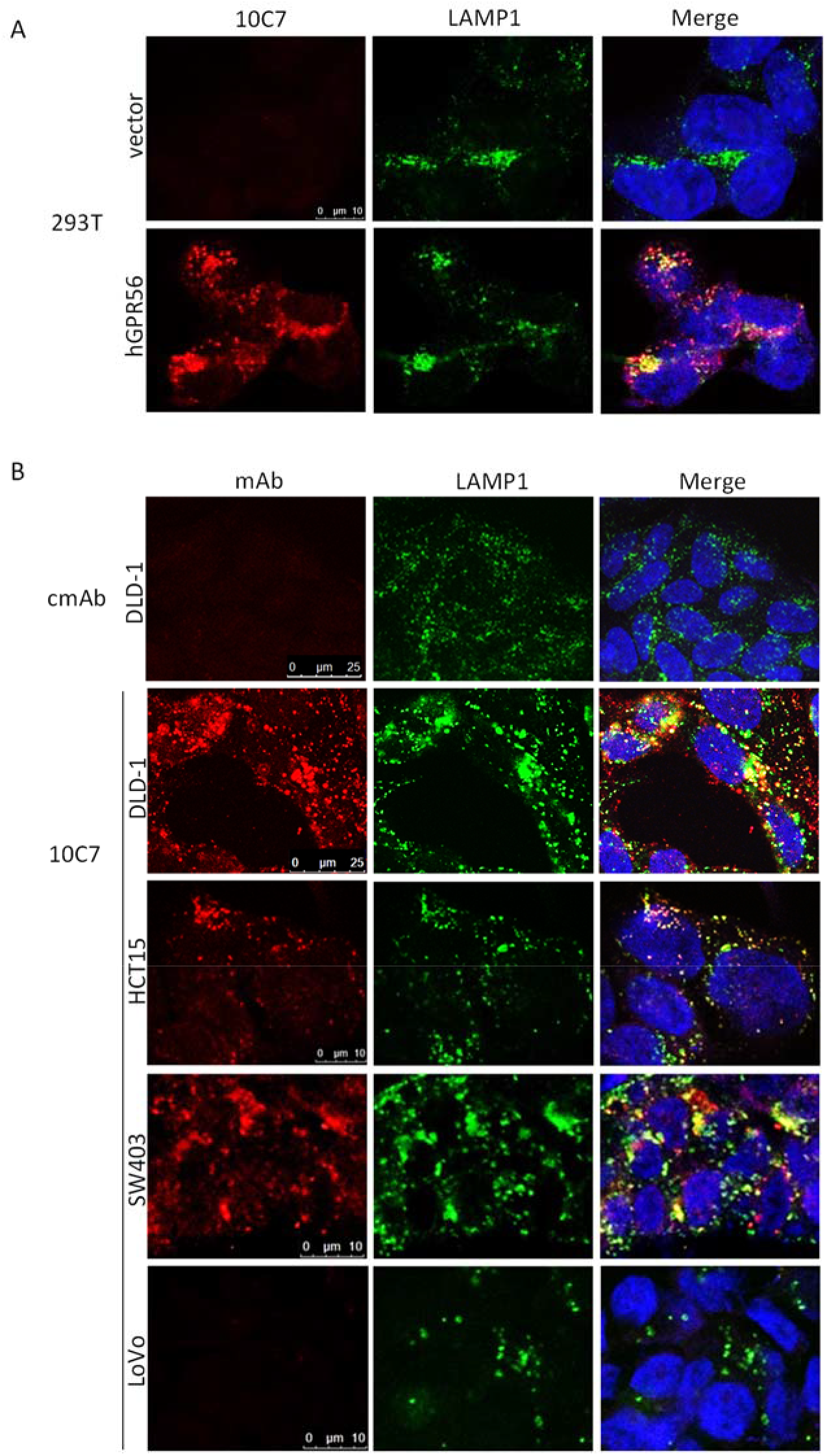
GPR56 mAb internalizes and traffics to the lysosomes in CRC cells. Confocal images show the complex of 10C7 bound to GPR56 co-localizes with lysosome marker, LAMP1, after 1 hour at 37°C in A, hGPR56, but not vector 293T cells and B, CRC cell lines DLD-1, HCT15, SW403, and LoVo. No binding of a non-targeting control mAb (cmAb) was detected nor was 10C7 in GPR56-negative LoVo cells. 293T and CRC cells were treated with 6.5 nmol/L and 30 nmol/L 10C7, respectively.

### Generation of a GPR56 ADC that maintains high affinity binding and specificity

To select an effective payload for development of a GPR56-targeted ADC, we screened CRC cell lines against a panel of payloads of different classes including microtubule inhibitors (MMAE, monomethyl auristatin E and DM1, mertansine), topoisomerase inhibitors (DX8951, exatecan), and a DNA-damaging agent (DMSA, duocarmycin SA). IC_50_ values were determined after 4 days of treatment with free drug by quantitative CellTiter-Glo assays (Fig. 4A). We show that CRC cells were more sensitive to the DNA minor groove-binding, alkylating agent duocarmycin (IC_50_ range = 93.6-696 pmol/L) compared to microtubule inhibitors (MMAE, IC_50_ range = 260-2973 pmol/L; DM1, IC_50_ range = 556-3170 pmol/L) and DX8951 (IC_50_ range = 367-690 pmol/L). We therefore generated a novel GPR56 ADC by coupling 10C7 with the most potent payload, duocarmycin SA, through a cleavable maleimide-containing peptide linker, mc-vc-PAB-DMEA(PEG2) (Fig. 4B). The linker-payload was conjugated to 10C7 via thiols of cysteine residues exposed under partial reduction conditions, as we have previously reported (19). A similar conjugation reaction was performed using nontargeted cmAb. As shown by Coomassie staining of SDS-PAGE, the heavy and light chains of the ADCs had higher molecular weights compared to unconjugated mAbs, suggesting successful conjugation of duocarmycin to 10C7 and cmAb (Fig. 4C and Supplementary Fig. S3). SEC HPLC confirmed minimal aggregation (≤ 2%) (Supplementary Figs. S4A). As determined by HIC, the conjugation reactions yielded ADC with near similar average drug-antibody ratios (GPR56 ADC, DAR = 3.54; Supplementary Fig. S4B and cADC, DAR = 3.9; data not shown), making them suitable for comparison of binding and functional efficacy. To confirm that payload conjugation did not affect binding, we performed a fluorescence-based binding assay using hGPR56 293T cells. As shown in Fig. 4D, both GPR56 ADC and mAb exhibited equivalent affinity and maximum binding. Dose-dependent GPR56 ADC-mediated cytotoxicity was then tested against vector, hGPR56 WT and H360S mutant 293T cell lines. GPR56 ADC induced potent cell-killing activity against WT cells (IC_50_ = 1.8 nmol/L) with no significant effect on vector or H360S cells (Fig. 4E), demonstrating high specificity and efficacy.

**Figure 4.**
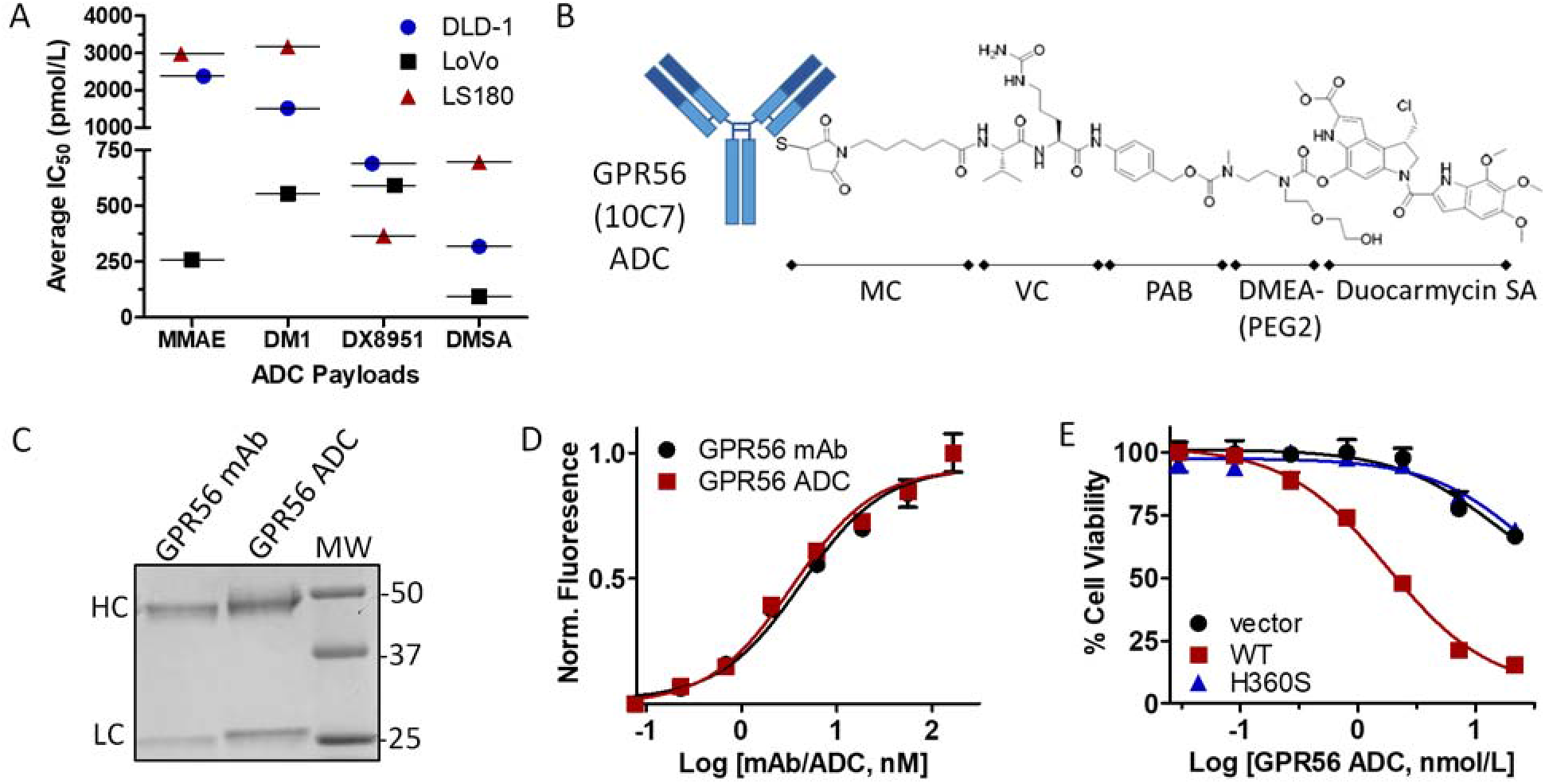
Generation and characterization of a GPR56-targeted antibody-drug conjugate. A, Plot showing IC_50_ values of ADC payloads for different CRC cell lines. MMAE, monomethyl auristatin; DM1, mertansine; DX8951, exatecan; DMSA, duocarmycin SA. B, Schematic depiction of the GPR56 ADC consisting of 10C7 mAb conjugated to the duocarmycin SA payload via a cleavable linker (DAR=3.54) attached to cysteine thiols. MC, maleimidocaproyl; VC, valine-citrulline; PAB, para-aminobenzyloxycarbonyl; DMEA, dimethylethanolamine; PEG, polyethylene glycol C, Coomassie stained SDS-PAGE of ~2μg GPR56 mAb compared to ADC under reducing conditions. Image shows shift in higher molecular weight (MW) of mAb after drug conjugation. HC, heavy chain; LC, light chain D, Cell-based binding assay confirms no change in binding of ADC compared to parent 10C7 mAb on hGPR56 293T cells. E, Cytotoxicity assay shows ADC selectively kills hGPR56 WT and not H360S mutant or vector 293T cells after 3 days. Experiments performed at least 3 times in triplicate. Error bars are SD.

### GPR56 ADC has potent cytotoxic activity against GPR56-high CRC cells and patient-derived tumor organoids

Next, we evaluated efficacy of GPR56 ADC against CRC cell lines expressing different levels of endogenous GPR56 (Fig. 1H). We first compared GPR56 ADC activity in DLD-1 cells with and without GPR56 shRNA knockdown (KD). GPR56 ADC had potent activity against control (CTL) shRNA cells (IC_50_ = 40 nmol/L) with minimal effect against GPR56 KD cells (i.e. only 40% reduction in viability at highest dose; Fig. 5A). Loss of GPR56 expression was confirmed by western blot (Fig. 5B). GPR56 ADC was then tested against GPR56-high CRC cell lines: SW620 (IC_50_ = 29.4 nmol/L), SW403 (IC_50_ = 3.7 nmol/L), HT-29 (IC_50_ = 9.1 nmol/L), and those with moderate levels of GPR56 expression: HCT15 (IC_50_ = 98 nmol/L), DLD-1 (parental, IC_50_ = 32.6 nmol/L), and LS180 (IC_50_ = ND; Fig. 5C-E). Though HCT15 and LS180 cells expressed moderate levels of GPR56, both cell lines were relatively insensitive to ADC-induced cytotoxicity. This may be attributed to intrinsic resistance to duocarmycin compared to other CRC cell lines, as shown for LS180 (Fig. 4A). GPR56 ADC showed relatively no effect on GPR56-low/negative CRC cell lines, RKO and LoVo, at the highest dose tested. No significant activity was measured by unconjugated GPR56 (10C7) mAb, nontargeted cmAb or cADC, as shown for SW620 and HCT15 cells (Fig. 5C-D). We next evaluated our GPR56 ADC against a patient-derived tumor organoid (PDO) model of metastatic rectal cancer (CRC-001). PDOs were shown to express high levels of GPR56 as confirmed by western blot (Fig. 5F). Lysate from hGPR56 293T and LS180 cells were loaded as positive controls for expression. PDOs were treated with 30 nmol/L GPR56 ADC, cADC, or vehicle (PBS) for 5 days. As shown in Fig. 5F-G, GPR56 ADC resulted in near complete tumor cell-killing, whereas cADC only reduced viability by approximately 30% (Fig. 5G-H). These results show that GPR56 ADC mediates target-specific cytotoxicity against GPR56-positve CRC cells and tumor organoids.

**Figure 5.**
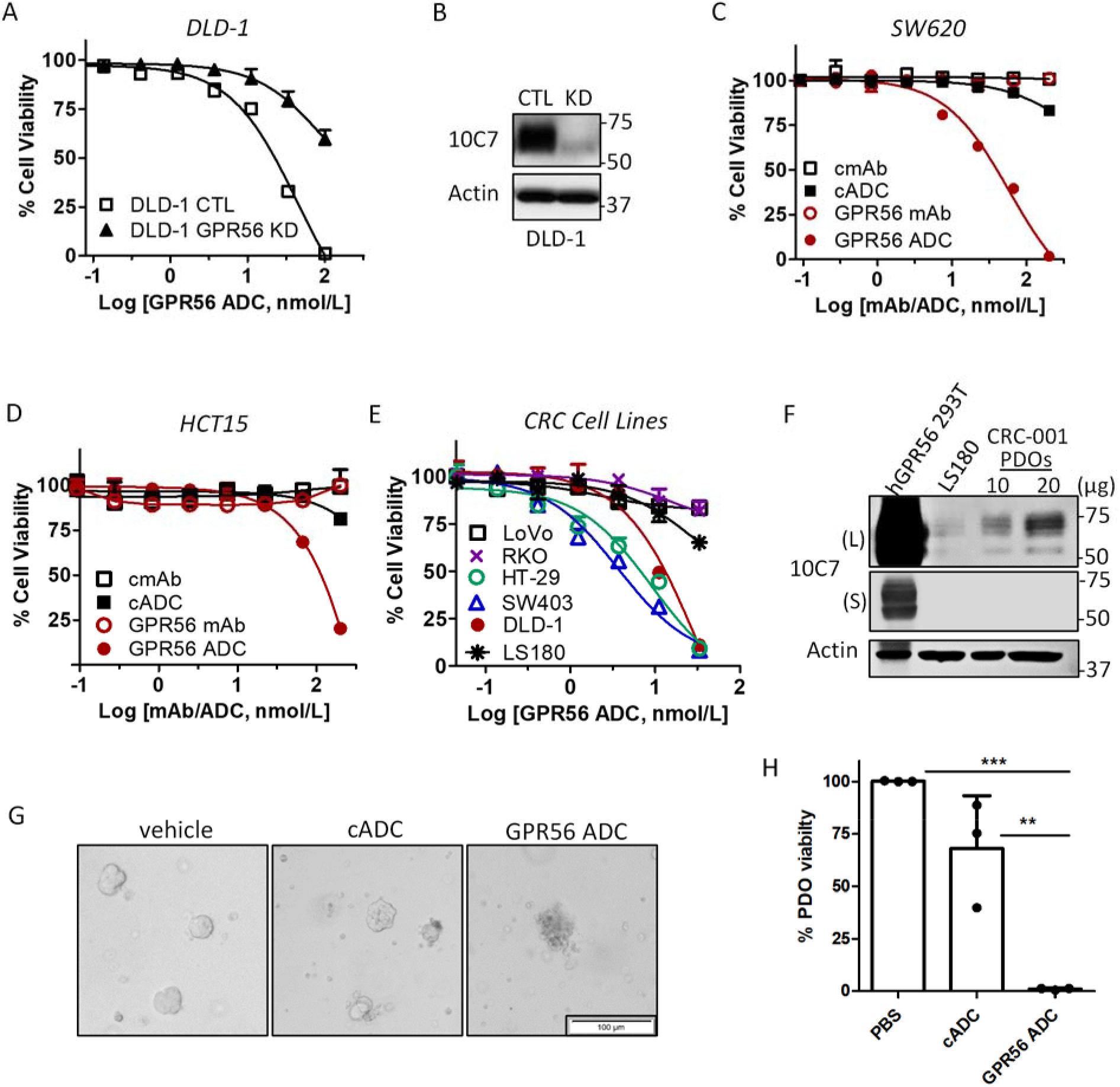
GPR56 ADC potently and selectively kills GPR56-high CRC cell lines and patient-derived tumor organoids. A, Cytotoxicity assay shows ADC eliminates DLD-1 control (CTL) cells, but not GPR56 knockdown (KD) cells, in a concentration dependent manner. B, Western blot of GPR56 expression in CTL and GPR56 shRNA DLD-1 stable cell lines. Cytotoxicity of GPR56 ADC against C, SW620 and D, HCT15 cells treated with GPR56 mAb, GPR56 ADC, non-targeting control mAb (cmAb), or control ADC (cADC). E, GPR56 ADC against multiple CRC cell lines with different levels of GPR56 expression. F, Western blot of GPR56 expression in patient-derived tumor organoids (PDOs) of a liver metastasis of rectal adenocarcinoma (CRC-001). Different amounts of PDO protein lysate was assessed. hGPR56 293T and LS180 lysates represent positive controls for GPR56 expression. L, long exposure; S, short exposure G, Representative brightfield microscopy images and H, cytotoxicity assay of PDOs treated with 30 nmol/L cADC, GPR56 ADC, or vehicle (PBS). Experiments were performed 2-3 times in triplicates. Statistical analysis was performed using oneway ANOVA. Error Bars are SEM.

### GPR56 ADC demonstrates antitumor efficacy in preclinical models of CRC

As 10C7 does not bind mGPR56, we were unable to test on-target ADC effects in genetic mouse models of CRC. However, a major issue in ADC development is off-target toxicity attributed to factors such as low stability and safety of the cytotoxic payload (24). We therefore conducted a single dose escalation study in immunocompetent C57BL/6 mice. Mice were administered vehicle (PBS), 4, 8, or 16 mg/kg GPR56 ADC and monitored for weight loss or other signs of overt toxicity. After 2 weeks, blood was collected and hematology and clinical chemistry analyses were performed. Mice showed no significant changes in weight, liver enzymes, or blood cell counts (Supplementary Fig. S5A-E). These results indicated that GPR56 ADCs were well-tolerated based on off-target assessment.

Anti-tumor efficacy was then evaluated against xenograft models of SW620, HT-29, and SW403 CRC cells (MSS subtype), which demonstrated highest potency in vitro. SW620 xenografts were dosed weekly with 1.5 and 5 mg/kg of GPR56 ADC. While 5 mg/kg GPR56 ADC was sufficient to induce slight tumor regression followed by stasis, 1.5 mg/kg only resulted in a reduction in tumor growth (~30%) after the third treatment. Treatment with unconjugated 10C7 mAb had no significant impact on tumor growth (Fig. 6A). The HT-29 model was treated with 2.5 and 5 mg/kg GPR56 ADC, resulting in a 20% and 40% reduction in tumor growth, respectively (Fig. 6B). Given that HT-29 tumors grew very rapidly, we were only able to dose once before the vehicle group had to be euthanized. However, we observed a substantial decrease in the rate of growth after a second dose, near tumor stasis, especially at the higher dosing level (Fig. 6B). GPR56 ADC tested at a dose of 2.5 mg/kg in SW403 xenografts showed a 50% reduction after 3 treatments (Fig. 6C). Treatment with nontargeted cADC had no significant impact on tumor growth (Fig. 6C). We next evaluated GPR56 ADC efficacy in two PDX models of CRC at the 2.5 mg/kg dose. We showed GPR56 ADC was effective against CRC-001 PDOs ex vivo (Fig. 5F-G) and, as shown by western blot, CRC-001 (liver metastasis of rectal cancer) and XST-GI-005 (colon cancer) MSS PDX tumors express high levels of GPR56 (Fig. 6D). GPR56 ADC treatment resulted in 70% and 50% inhibition of tumor growth in CRC-001 and XST-GI-005 models, respectively (Fig. 6E-F). No significant effect in body weight or overt toxicity was observed during treatment in the nu/nu or NSG mouse strains (Fig. 6G-H). These findings demonstrate GPR56 ADC has significant efficacy against GPR56-expressing CRC tumors in vivo.

**Figure 6.**
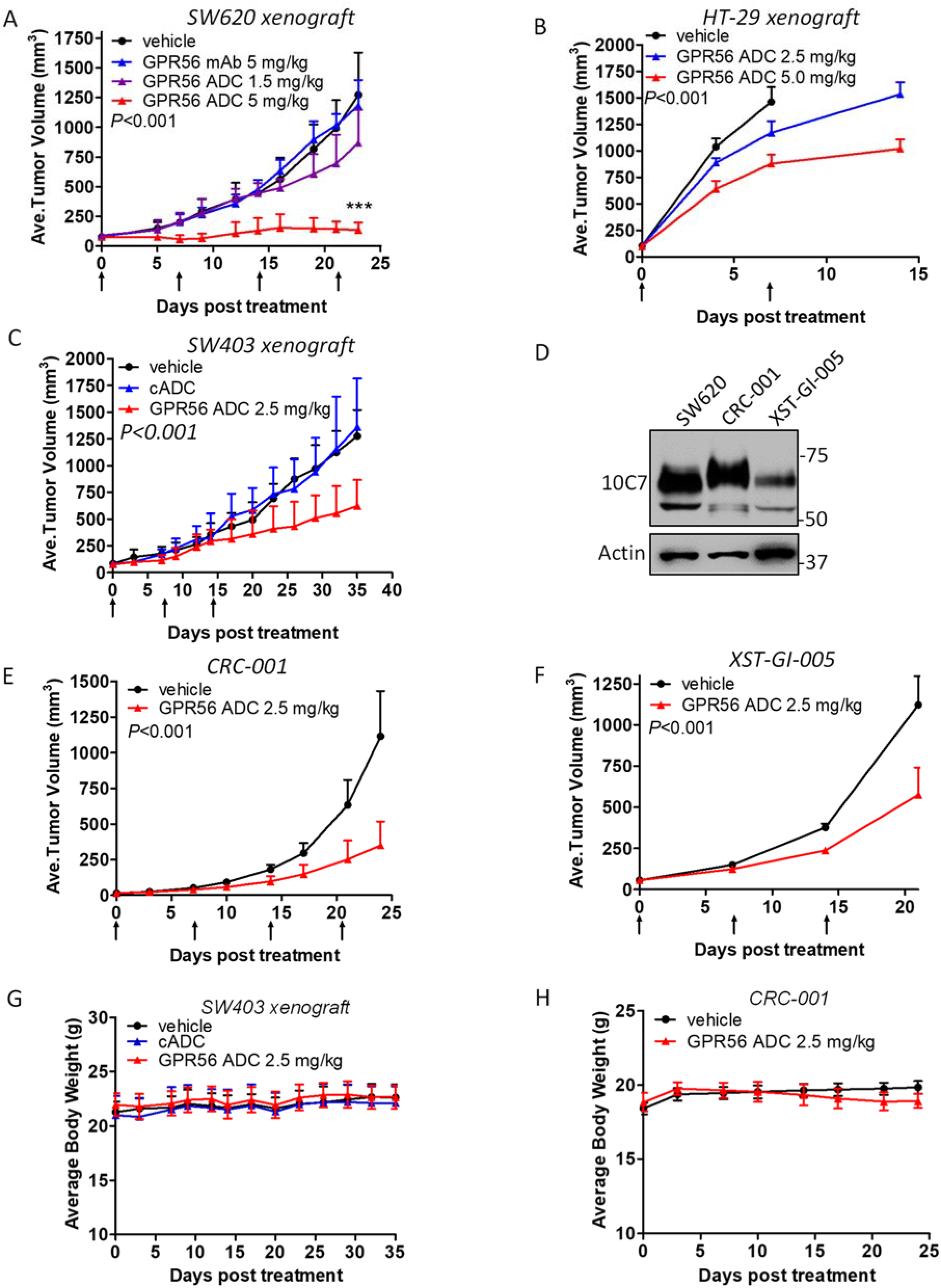
GPR56 ADC induces antitumor activity CRC cell line and patient-derived xenograft models. In vivo efficacy of GPR56 ADC was evaluated in A, SW620 (n=6, vehicle; n=4, GPR56 mAb and 1.5mg/kg ADC; n=5, 5 mg/kg ADC), B, HT-29 (n=5/group), and C, SW403 (n=6, vehicle; n=5 cADC and GPR56 ADC) xenograft models. D, Western blot of GPR56 expression in PDX models of CRC. SW620 lysate was used as a positive control. In vivo efficacy of GPR56 ADC in PDX models E, CRC-001 (n=6) and F, XST-GI-005 (n=5). Bodyweight measurements of G, SW403 (nu/nu) and H, CRC-001 (NSG) tumor-bearing mice. Statistical analysis was performed using Student’s t-test for two groups or one-way ANOVA and Dunnett’s multiple comparison test. Error bars are SD.

## Discussion

Despite the availability of various therapeutic options for CRC treatment, the outlook for patients with advanced disease, especially the MSS subtype, remains poor. Chemotherapy is associated with many adverse side-effects and limited response and response to targeted therapies, such as cetuximab and panitumumab, is largely dependent on mutation statuses (2,25). Moreover, resistance to these therapies remains a major impasse for low survival rates. More recently, MSI-H status has emerged as a predictor of remarkable sensitivity to immune checkpoint inhibitors (26). Though immunotherapy has significantly improved treatment outcomes for ~5% of MSI-H patients, a significant benefit has not been shown for those with MSS (25). In effect, patients with MSI-H tumors have better prognosis compared to patients with MSI-L/MSS tumors (26), thus signifying the urgent need for new therapies to treat MSI-L/MSS tumors. We showed GPR56 correlates with poor prognosis in CRC (11) and its expression is highest in tumors that are MSI-L/MSS and CIMP-L/non-CIMP (Fig. 1). The finding that a large percentage of CRC tumors are categorized within these molecular subtypes suggests that a GPR56-targeted ADC has the potential to treat a considerable patient population. Though *GPR56* expression negatively correlated with mutation in *BRAF* (Fig. 1G), due to *BRAF* association with MSI-H/CIMP-H status, a GPR56 ADC would likely be effective irrespective of other common mutations associated with CRC (i.e. *APC, KRAS, TP53*).

In ADC development, selection of an appropriate payload is essential. Payloads, depending on their mechanism of action, can have different efficacy against different cancer types. For example, microtubule inhibitors have been shown to be generally ineffective against CRC and other GI cancers (27), consistent with our data (Fig. 4A). LoVo cells are an exception, as they are more sensitive to MMAE and DM1 compared to other CRC cell lines. Multidrug resistance (MDR) is another important factor that can hinder the therapeutic effect of ADCs in the cancer treatment. Interestingly, we previously showed that GPR56 enhanced drug resistance in CRC cells through upregulation of the drug efflux pump, P-glycoprotein (P-gp)/MDR1 (11). We also found that treatment of MMAE-resistant CRC cells with tariquidar, a P-gp/MDR1 inhibitor, partially sensitized them to an MMAE-conjugated LGR5-targeted ADC (11). Duocarmycins, on the other hand, are DNA-alkylating agents that have the inherent capacity to better evade traditional resistance mechanisms and retain high potency against MDR cancer cells (28,29). Duocarmycin-based ADCs that target HER2 (SYD985) and B7-H3 (MGC018) are currently being evaluated in clinical trials for the treatment of solid tumors, with several more at preclinical stages (30,31).

We show that a duocarmycin-based GPR56 ADC is highly potent at inducing cytotoxicity in CRC. In vitro studies demonstrate high affinity binding, effective internalization, and selective delivery of payload to CRC cells and tumor organoids with high GPR56 expression (Figs. 2-5). Significant antitumor activity was shown across several CRC cell line xenografts and PDX models (Fig. 6). As optimal ADC dosing concentrations for humans are typically much lower than 5mg/kg (32), concentrations beyond this level were not evaluated for this study. Future work to improve ADC efficacy will include affinity maturation, increasing the DAR, and conjugation to more potent payloads, such as a pyrrolobenzodiazepine that has been used to generate the recently approved ADC, loncastuximab tesirine (33).

Our preliminary off-target safety study showed no obvious signs of toxicity. However, for the successful development of an ADC both off-target and on-target effects need to be evaluated. Analyses of normal tissues showed highest GPR56 expression in mouse and human in areas of the brain, kidney, pancreas, and thyroid (human), though at lower levels compared to tumors (10,13–15), and consistent with TCGA datasets (12). We do not anticipate that a GPR56-targeted ADC would have adverse effects on the brain as mAbs generally do not cross the blood-brain barrier (BBB). The BBB constitutes tight junctions between endothelial cells, restricting the influx of large molecules such as mAbs (34). Interestingly, the FDA-approved ADC, sacituzumab govitecan, is directed against the target TROP-2 which has relatively high expression in many normal tissues (12,35). Therefore, to appropriately address potential systemic on-target effects, future testing of a surrogate mAb that binds GPR56 in rodent models is needed. GPR56 expression has also been detected on human natural killer (NK) cells, and shown to inhibit effector functions during their inactive state (36,37). How relative surface levels of GPR56 on NK cells compares to tumor cells and the impact of a GPR56 ADC on the immune system needs to be further examined.

The GPR56 ECD is structurally characterized by a highly conserved GAIN domain containing a juxtamembrane GPS that can be autoproteolytically cleaved, leaving two noncovalently associated but distinct fragments, the N-terminal fragment (NTF) and C-terminal fragment (CTF) or stalk region (38,39). We showed that 10C7 binds the ECD within the GAIN domain of the NTF and potentiates GPR56-mediated Src-Fak and SRF-RE signaling in vitro (18). Here, we further pinpointed the epitope and identified H360, proximal to the GPS site (L382/T383), as an essential residue for 10C7 binding and activation of hGPR56 (Fig. 2). This reveals an important region for inducing a conformational change in the NTF that results in GPR56 activation and a potential site for interaction(s) with cognate ligands or co-receptors. This finding opens doors for further investigation of the GPR56 signaling mechanism in normal cells and cancer.

In summary, we have identified GPR56 as a novel target for ADC development. These preclinical data provide proof-of-concept that targeting GPR56 using the ADC approach may be an effective strategy for the treatment of GPR56-expressing colorectal tumors, especially those 80-85% classified as MSI-L/MSS. As GPR56 is also highly expressed in acute myeloid leukemia, glioblastoma, melanoma and tumors of the ovary, breast, lung, and pancreas (8,9,12,15,40,41), a GPR56-targeted therapy may have broad utility for the treatment of many cancer types.

## Authors’ Contributions

Conceptualization and Design: K.S.C; Methodology: K.S.C.; Investigation: K.S.C., L.E.F., J.J., T.C., S.S., Z.L., Q.J.L., J.R.; Data curation: K.S.C., J.J., L.E.F., T.C., Z.L., S.S.; Formal Analysis: K.S.C., L.E.F., J. J., T.C., Z.L., S.S.; Writing original draft: K.S.C. and J.J.; Review and Editing: K.S.C, J.J., Q.J.L.; Resources: K.S.C., Q.J.L., J.R.; Funding acquisition: K.S.C. and Q.J.L.; Supervision: K.S.C.

## Acknowledgements

This work was supported by funding from the National Institutes of Health (NCI R01 CA226894) and Welch Foundation Endowment Fund Award (L-AU-0002-19940421) to K.S.C., funding from the Cancer Prevention and Research Institute of Texas (RP190542) to Q.J.L. and K.S.C., a fellowship of the Gulf Coast Consortia, on the Training Interdisciplinary Pharmacology Scientists Program (T32 GM139801) to J. J., and a Cancer Therapeutics Training Program Fellowship (RP210043) to L.E.F. We would like to thank Sheng Zhang and Wangsheng Yu for technical contributions, Betty Arceneaux and Dr. Karan Saluja with assistance in patient sample collection and processing, and Martha Thompson for assistance with regulatory approvals for research involving human subjects.

