## Supplementary Figures S1-S5 and Supplementary Table 1 for "An antibody-drug conjugate targeting GPR56 demonstrates efficacy in preclinical models of colorectal cancer"

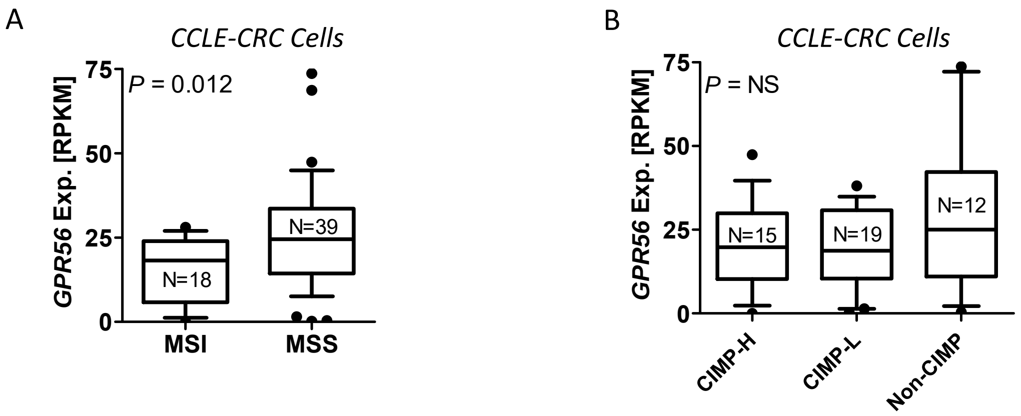


**Supplementary Figure S1. GPR56 expression in CRC cells of different subtypes.** Distribution of A, MSS/MSI and B, CIMP subtypes based on GPR56 expression analysis of CRC cell lines from the Cancer Cell Line Encyclopedia (CCLE).

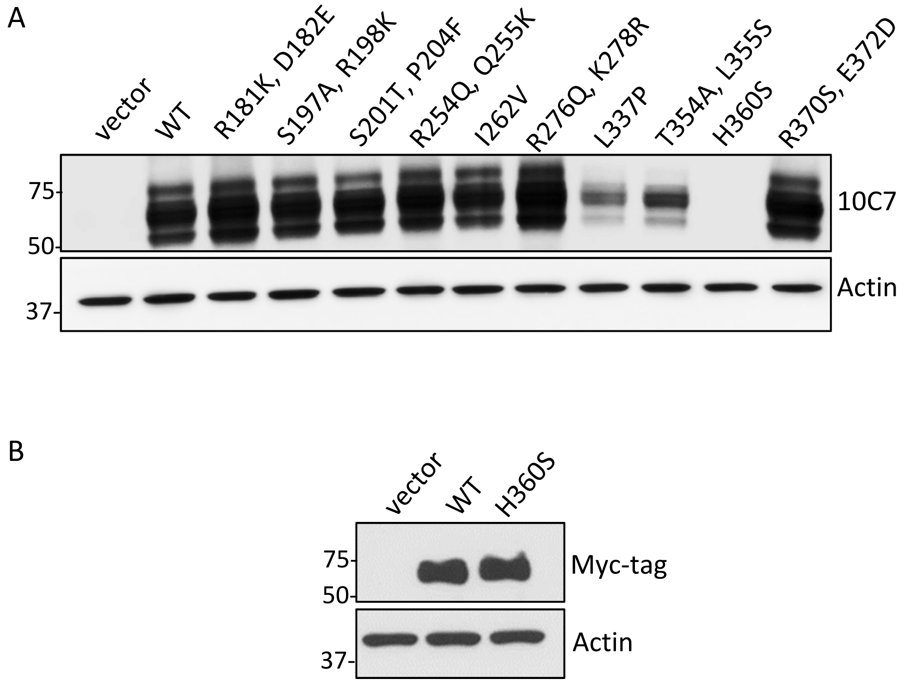


**Supplementary Figure S2. 10C7 epitope mapping.** Western blots of A, 10C7 detection of hGPR56 WT and different mutants and B, confirmation of equivalent expression of WT and H360S mutant for SRF-RE assay.

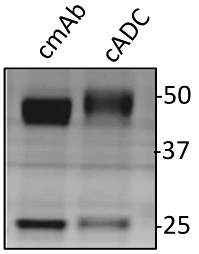


**Supplemental Figure S3. Nontargeted control mAb and ADC.** Coomassie stained SDS-PAGE of ~3µg nontargeted control hIgG1 mAb (cmAb) compared to control ADC (cADC) under reducing conditions. Image shows shift in higher molecular weight (MW) of mAb after drug conjugation. HC, heavy chain; LC, light chain

A

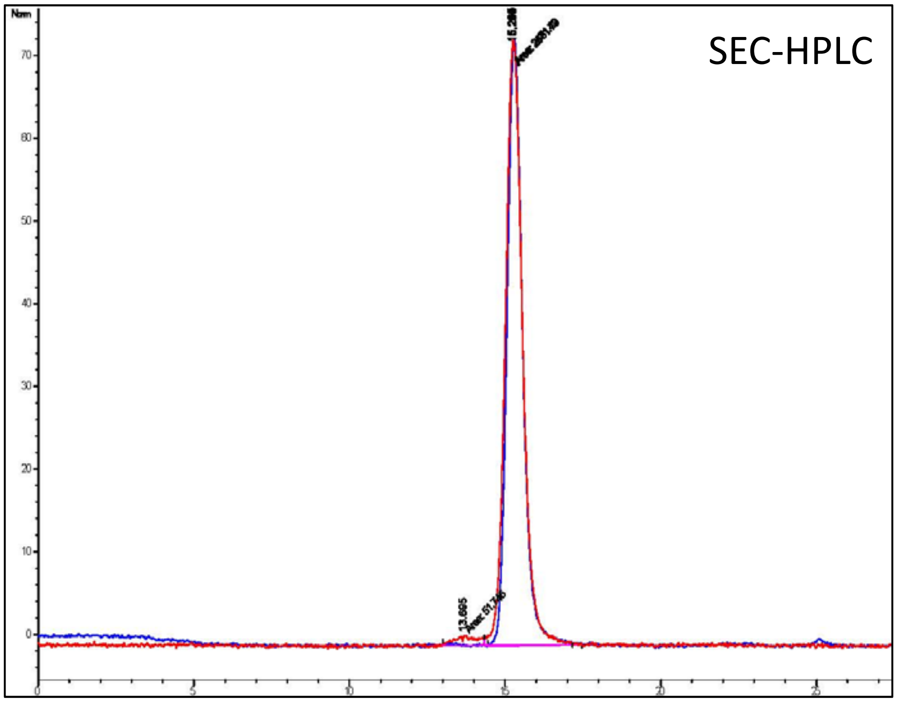


B

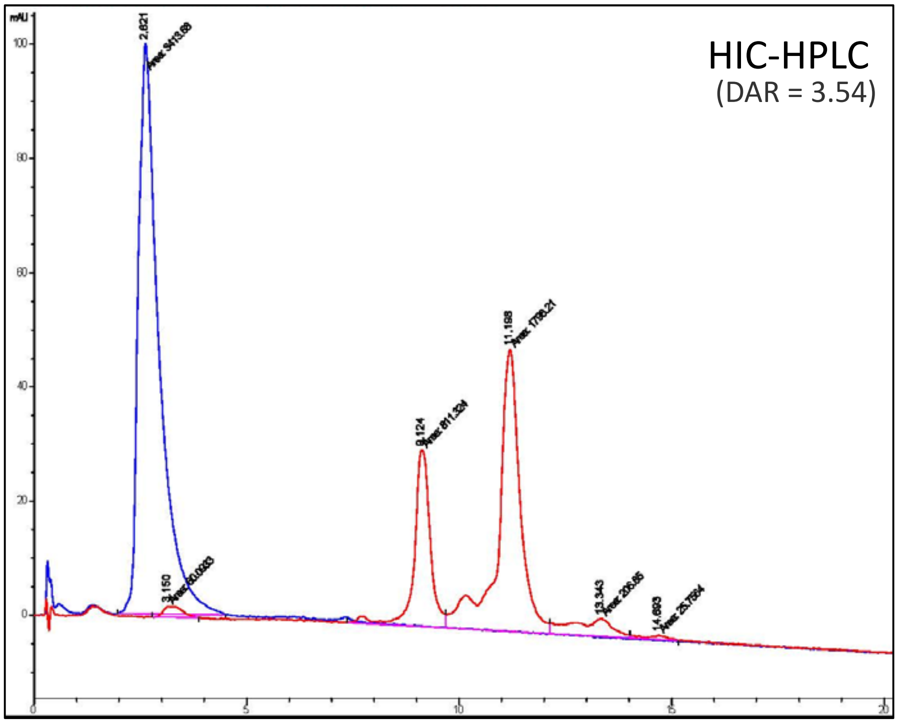


**Supplementary Figure S4. Analytical method profiles for GPR56 ADC and its corresponding 10C7 mAb. A,** Size Exclusion Chromatography (SEC) shows ADC (red) and mAb (blue) monomers with ~2% ADC aggregation. **B**, Hydrophobic Interaction Chromatography (HIC) shows unconjugated mAb and ADC (left side peaks) and ADC conjugated with different number of drug molecules (red peaks). Average DAR = 3.54.

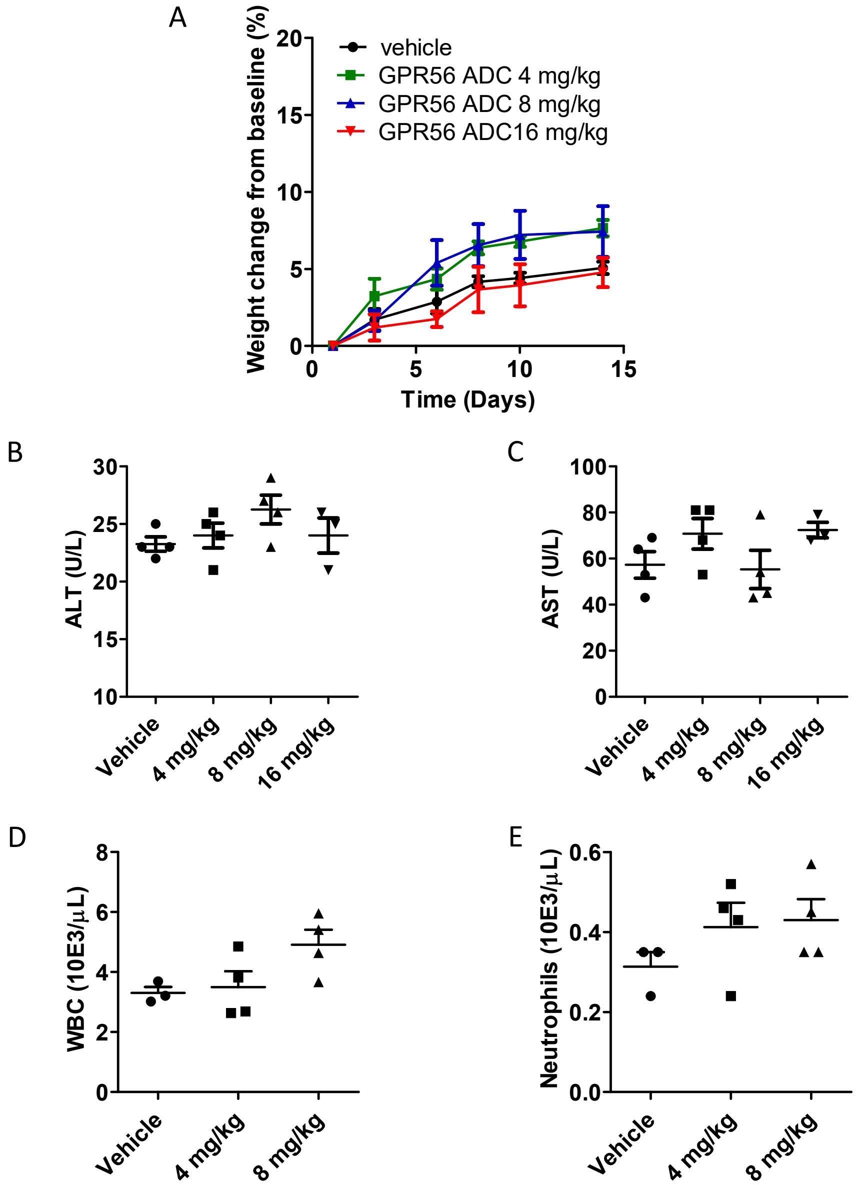


**Supplemental Figure S5. Off-target safety assessment of GPR56 ADC.** Immunocompetent C57/BL/6 mice were treated on day 1 with single dose GPR56 ADC, as indicated, or vehicle. Assessment of Clinical chemistry and hematology was performed in the Department of Veterinary Medicine & Surgery at MD Anderson Cancer Center. A, Bodyweights of mice (n=4 for each group except 16mg/kg, n=3). B, Alanine aminotransferase (ALT) and C, aspartate aminotransferase (AST) liver enzyme analysis on serum collected from mice on day 14 (n=4 for each group except 16mg/kg, n=3). D, White blood cell (WBC) and E, neutrophil counts on day 14 (vehicle, n=3; GPR56 ADC 4 mg/kg and 8mg/kg, n=4).

**Supplementary Table 1. Mutagenesis Primers**

| GPR56 Mutant (Human→ Mouse) | Forward Primer (5'-3') | Reverse Primer (5'-3') |
|---|---|---|
| R181K, D182E | ATGTGCGAGCTCAAAAAGGAACTCCAGCTGCTCAG | CTGGCTGAGCAGCTGGAGTTCCTTTTTGAGCTCGCAC |
| S197A, R198K | ATCCCCAGAAGGCCGCAAAGAGGCCCTCGGCTG | CAGCCGAGGGCCTCTTTGCGGCCTTCTGGGGAT |
| S201T, P204F | TCAAGGAGGCCCACGGCTGCCTTCGCCAGCCAG | CTGGCTGGCGAAGGCAGCCGTGGGCCTCCTTGA |
| R254Q, Q255K | ACATCCACTCCCAGAAGGAGGAGGAGCAGAG | CTCTGCTCCTCCTCCTTCTGGGAGTGGATGT |
| I262V | AGGAGCAGAGCGAGGTCATGGAGTACTCGGTGC | GCACCGAGTACTCCATGACCTCGCTCTGCTCCT |
| R276Q, K278R | TCGAACACTCTTCCAGCAGACGAGAGGCCGGAGCGG | CCGCTCCGGCCTCTCGTCTGCTGGAAGAGTGTTCGA |
| L337P | TCACCTTCCAGCACCAGCCACAGCCGAAGAATGTGAC | GTCACATTCTTCGGCTGTGGCTGGTGCTGGAAGGTGA |
| T354, L355S | GTTGAAGACCCCGCATCGAGCAGCCCG | CGGGCTGCTCGATGCGGGGTCTTCAAC |
| P358T | CCACATTGAGCAGCACGGGGCATTGGAGCAGTGCTG | CAGCACTGCTCCAATGCCCCGTGCTGCTCAATGTGG |
| H360S | GAGCAGCCCGGGGTCTTGGAGCAGTGCTGGGTGT | ACACCCAGCACTGCTCCAAGACCCCGGGCTGCTC |
| R370S, E372D | GGTGTGAGACCGTCAGCAGAGATACCCAAACATCCTGC | GCAGGATGTTTGGGTATCTCTGCTGACGGTCTCACACC |
